# Buffering of genetic defects in animal development by regeneration programs

**DOI:** 10.1101/2025.10.25.684558

**Authors:** Kazunori Ando, Sushant Bangru, John Welsby, John D. Thompson, Kenneth D. Poss

## Abstract

Regeneration programs enable animals to restore damaged or lost tissues, and the range of stimuli for these programs is incompletely understood. Here, we used zebrafish, a vertebrate species with exceptional regenerative capacity, to identify chemically induced mutations that alter regeneration-associated gene activation. Transgenic zebrafish with a permissive promoter and EGFP cassette inserted in the vicinity of the pro-regenerative factor gene *fgf20a* were mutagenized, and larvae homozygous for ENU-induced mutations were assessed for disruptions in *fgf20a*-directed reporter gene expression following fin fold amputation. One line was identified with heritable, elevated *fgf20a*:EGFP presence in the absence of experimental injury, localized to regions of fin fold tissue undergoing degeneration. Whole-genome sequencing (WGS) identified a mutation within exon 72 of the *fraser syndrome 1* (*fras1*) gene, mutated in patients with inherited skin disease. *fras1* mutant larvae spontaneously displayed elevated expression of other known injury/regeneration-responsive reporter lines in fin fold, and zebrafish crispants for homologs of other genes mutated in human developmental diseases also displayed regeneration-associated gene expression in regions of dysmorphology. Tempering Fgf signaling by transgenic expression of a dominant-negative Fgf receptor in *fras1* mutants exacerbated the disease phenotype. Our findings provide evidence that regeneration programs are harnessed in response to developmental defects caused by genetic mutations, potentially buffering deleterious phenotypes.

## INTRODUCTION

One of the most striking biological disparities between vertebrate species is their varied capacity to regenerate damaged tissues. Whereas in many adult mammalian tissues the regenerative response to injury is limited, involving fibrosis and incomplete repair, organisms like salamanders and zebrafish can regenerate fins, limbs, heart, spinal cord, kidney, and other tissues/organs with remarkable fidelity^1,2^. Across these diverse tissue contexts, regeneration relies on coordinated activation of transcriptional programs that trigger cell proliferation, establish tissue architecture and restore tissue function. In recent years it has become clear that regeneration programs are orchestrated by dedicated *cis*-regulatory elements^3,4^. First referred to as tissue regeneration enhancer elements (TREEs), and called regeneration responsive elements (RREs), these sequences integrate injury signals with developmental gene networks to selectively activate transcription in damaged tissue. For example, the Fgf ligand gene *fgf20a* is induced within hours of fin amputation in zebrafish, where it is required to initiate formation of the regeneration blastema^5,6^, several TREEs appear to collectively direct its contextual expression^7^. Enhancers near the *leptinb, il11a, inhbaa, ven* genes similarly direct target gene transcription after fin amputation^3,8,9^. Many other TREEs have been identified across diverse species, tissues, cells, and injury contexts^10–13^.

A wide range of injury-associated cues can trigger regeneration programs. Reactive oxygen species, particularly hydrogen peroxide, form tissue-scale gradients within minutes of wounding and recruit leukocytes to initiate the regenerative response^14^. Innate immune activity is equally critical: macrophage-derived cytokines regulate blastema formation and patterning in both salamander limbs and zebrafish fins^15,16^. Oxygen tension provides another axis of control, as hypoxia stimulates cardiomyocyte dedifferentiation and proliferation in zebrafish hearts and influences regeneration in mammals^17,18^. Hormones are thought to gate regenerative competence; thyroid hormone suppresses cardiac regeneration, whereas disruption of thyroid hormone receptor signaling can enhance repair by altering metabolic and hypoxia pathways^19–21^. Innervation, and mechanical and hydrodynamic strain, are also early influences on regeneration^22,23^. Each of these triggering signals ultimately converge on regulatory controls like enhancers.

Fluctuations in signals with impact during regeneration also occur during normal development and tissue homeostasis. Tissue growth, turnover, and morphogenesis naturally involve episodes of hypoxia, mechanical strain, inflammation, and epithelial remodeling. The zebrafish heart, for example, undergoes a period of strain during the juvenile stage that initiates a program with several commonalities as injury-induced heart regeneration in adults^24^. During lung branching morphogenesis, rhythmic tissue expansion generates epithelial tension that activates YAP signaling to drive proliferation and epithelial remodeling— mechanisms like those redeployed during adult lung repair and regeneration^25^. Likewise, physiological hypoxia serves as a morphogenetic cue during early embryogenesis, where low oxygen tension enhances WNT pathway activity and primitive-streak gene expression through HIF-dependent regulation—mirroring hypoxia’s role in promoting tissue growth and angiogenesis during regeneration^26^. Thus, it is likely that regeneration programs may act as general surveillance systems, engaged at any point in life when tissues experience stress or compromised integrity—even during development.

Here, we conducted a forward genetic screen using a zebrafish *fgf20a* enhancer trap line to identify mutations that disrupt activation of regeneration-associated gene expression upon amputation injury in fins. Unexpectedly, we identified a line harboring recessive mutations in *fras1*, a gene previously linked to epithelial integrity and extracellular matrix organization in many organisms from zebrafish to humans, that caused spontaneous activation of *fgf20a* regulatory sequences. To explore a hypothesis that enhancer-driven regenerative networks are deployed in response to intrinsic developmental defects, we analyzed additional regeneration-responsive gene expression in these mutants, performed crispant analysis of several other genes associated with human developmental disorders, and experimentally limited Fgf signaling during *fras1* deficiency. Our results suggest a new and potentially important role for injury-induced gene expression programs as protective systems to buffer the effects of deleterious gene mutations.

## RESULTS

### ENU mutagenesis screen for mutations that alter regeneration-associated *fgf20a* expression

Enhancer trap and reporter lines for injury-responsive genes enable real-time visualization of regeneration-associated gene expression and can be powerful tools for dissecting the molecular logic of regeneration (**Fig. 1A**). To identify potential regulatory factors required for an essential factor for fin regeneration, *fgf20a*, we performed ENU mutagenesis in homozygous males of an *fgf20a* enhancer trap line (*fgf20a*:*EGFP*; **Fig. 1B**)^27^. In this line, a transposon-based *hsp70l*:*EGFP* cassette randomly inserted 1.5 kb upstream of the *fgf20a* gene locus. *fgf20a*:*EGFP* animals display strong, injury-induced EGFP expression after caudal fin amputation or larval fin fold amputation, mimicking endogenous *fgf20a* expression^28^. We treated 11 homozygous *fgf20a*:*EGFP* male fish (F_0_) with ENU, and, after confirming a high mutagenesis rate, we prepared 117 F_2_ families, all homozygous for *fgf20a*:*EGFP*, to breeding age by intercrossing 190 F_1_ fish **(Fig. 1C**). Larval fin fold regeneration is robust in zebrafish, and both *fgf20a* mRNA and *fgf20a*:*EGFP* are induced within one day of amputation of the 3 dpf larval fin fold **(Fig. 1D**)^6^. Thus, we screened 9868 total animals from 460 F3 families (all homozygous for *fgf20a*:*EGFP* and for ENU-induced mutations) for induction of *fgf20a*:*EGFP* after larval fin fold amputation.

**Fig. 1.**
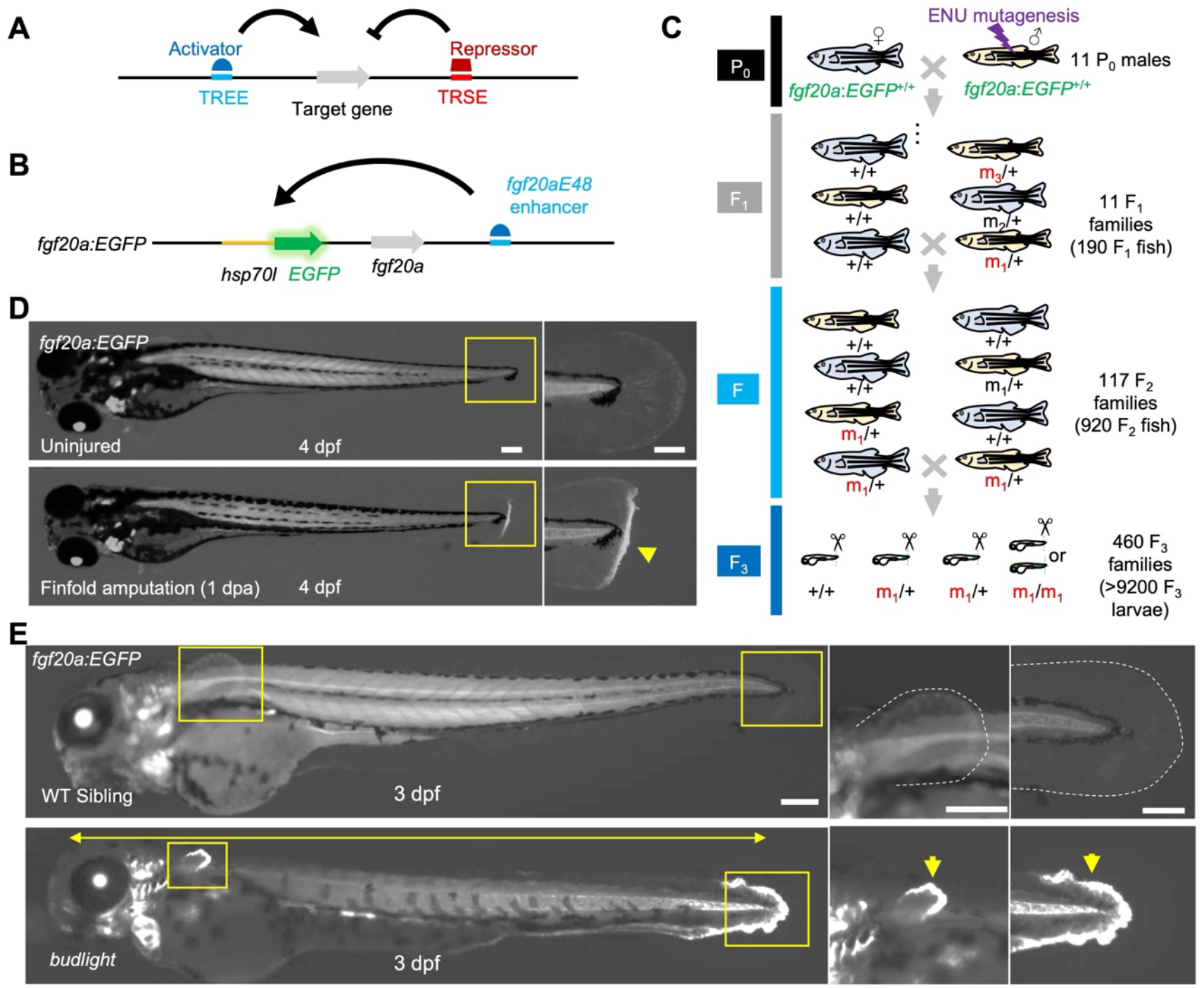
ENU mutagenesis identified a mutation that increases *fgf20a*-directed expression at the degenerating finfold. **(A)** Regeneration-responsive genes are regulated by tissue regeneration enhancer elements (TREEs) and tissue regeneration silencer elements (TRSEs). **(B)** Regeneration-responsive expression of the pro-regenerative factor, *fgf20a*, is regulated by a TREE, *fgf20aE48*, also occurring in the enhancer trap transgenic reporter line, *HGn21A*. **(C)** ENU mutagenesis screen flow. Homozygous male *fgf20a:EGFP* fish were treated with ENU and mated with homozygous *fgf20a:EGFP* females for F_1_ families. F_2_ families were generated from intercrossing F_1_ fish, and screens of F_3_ larvae were performed to detect homozygous mutations that reduce or increase injury-induced fluorescence (m_1_, m_2_, m_3_, …). **(D) (Top)** Uninjured 4 dpf larva. **(Bottom)** 4 dpf larva after finfold amputation at 3 dpf. EGFP expression is clearly induced at the injury site (yellow arrowhead in the right magnified view of tail region), mimicking fgf20a induction after the same injury. **(E)** The heritable mutant larvae (*budlight*) identified from the screens spontaneously had short body phenotype (yellow double-headed arrow) and displayed an elevated expression of fgf20a:EGFP in the degenerated fin fold (yellow arrows).

From these 460 families, 144 contained larvae with potential changes in *fgf20a:EGFP* reporter expression and/or defects in regeneration. In reassessments of these families, 11 displayed abnormal regeneration 1 day after fin fold amputation. The majority of families contained larvae displaying limited fin fold regeneration, but also with developmental defects like finfold degeneration, cardiac edema, craniofacial malformation, or unusual trunk curvature **(Fig. S1)**. Two of these families had members displaying subtle defects in fin regeneration at the adult stage -- apparent delays in initiating regeneration but recovery to normal regeneration **(Fig. S2)**. Interestingly, while several families had increased *fgf20a:*EGFP in finfold or trunk muscle, all of these families showed roughly normal *fgf20a:EGFP* expression at the amputation site. Only one of the families had members displaying a gross change in fin fold *fgf20a:*EGFP from clutchmates. These larvae displayed a jagged, severely degenerated fin fold in the absence of injury, along with a conspicuous increase in *fgf20a:*EGFP at the caudal and ventral fin folds, and in pectoral fin buds, and thus we referred initially to this mutant as *budlight* (**Fig. 1E)**. In summary, ENU mutagenesis revealed a heritable mutation that causes spontaneous elevation of the *fgf20a* regeneration program in uninjured fin primordia undergoing degeneration.

### A nonsense mutation in *fras1* causes ectopic *fgf20a*-directed gene expression

To identify the molecular basis of *budlight*, we performed whole-genome sequencing using genomic DNA from 5 dpf wild-type larvae homozygous for the *fgf20a:EGFP* insertion, as well as from *budlight* siblings displaying the degenerating fin fold phenotype **(Fig. 2A)**. Sequence reads were aligned to the zebrafish reference genome, and we applied filtering steps to remove background variants, sequencing artifacts, and low-quality calls. This filtering enriched for homozygous variants unique to the mutant pool, thereby prioritizing candidate lesions most likely to underlie the phenotype. SNP homozygosity analysis revealed a single point mutation in exon 72 of the *fraser extracellular matrix complex subunit 1* (*fras1*) gene, corresponding to the third-to-last exon. This variant represented a nonsense mutation (T to A), introducing a premature stop codon in place of tyrosine at amino acid position 3708 **(Fig. 2B)**.

**Fig. 2.**
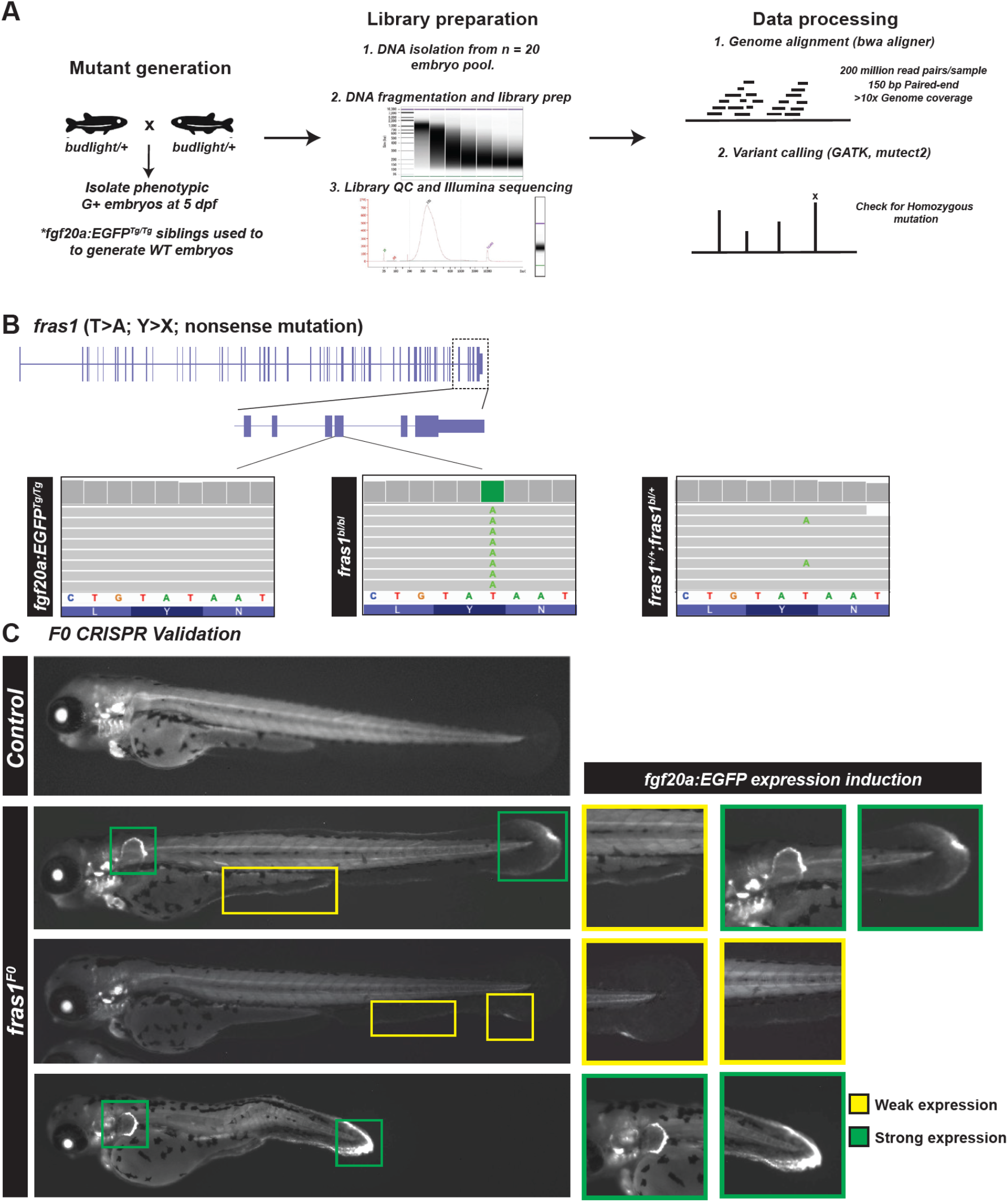
Mutants with elevated *fgf20a*:EGFP harbor a nonsense mutation in *fras1*. **(A)** Workflow schematic for whole genome sequencing of homozygous mutant, and HGn21 transgene homozygous controls. **(B)** WGS identified a single point mutation in the 72^nd^ exon of *fras1*, leading to a T to A conversion that results in replacement of a tyrosine (TAT codon) amino acid with a premature stop codon (TAA). **(C)** CRISPR-Cas9 F_0_ screening was used to validate *fras1* as the gene underlying the phenotype observed in the *budlight* ENU mutant.

*fras1* encodes a transmembrane protein with domains including von Willebrand factor–C and furin-like repeats, and it is crucial for epithelial–mesenchymal adhesion and basement membrane stability. In humans, mutations in FRAS1 cause Fraser syndrome, an autosomal recessive developmental disorder marked by epidermal blistering, syndactyly, and renal anomalies^29^. In zebrafish, *fras1* is expressed in epithelial tissues, particularly in developing fin folds, and is required for skin integrity and craniofacial morphogenesis^30^. Additionally, zebrafish mutants in *fras1*, as well as orthologues of FREM1/2 and other components of the Fraser syndrome complex, display classic fin blistering phenotypes that recapitulate aspects of the human and mouse diseases^31^.

To independently test whether *fras1* disruption was responsible for the *budlight* degenerative phenotype and activation of the *fgf20a:EGFP* enhancer trap, we designed CRISPR/Cas9 guides targeting *fras1* coding sequences. Injection of gRNAs into one-cell stage embryos generated mosaic F_0_ animals that phenocopied the ENU mutant: at 3 dpf, injected embryos exhibited localized fin fold degeneration accompanied by ectopic *fgf20a* induction in the pectoral fin bud, ventral fin fold, and caudal fin fold **(Fig. 2C)**. These CRISPR experiments establish a causal link between *fras1* loss-of-function and the spontaneous activation of regeneration programs observed in *budlight* mutants, which we refer to hereafter as *fras1* mutants.

### Mutants in zebrafish orthologues of human congenital disorder genes show ectopic *fgf20a*-directed gene expression

Based on the association of *fras1* mutations with *fgf20a* induction, we postulated that, more broadly, regeneration programs can be triggered by intrinsic defects in tissue architecture that arise from genetic lesions in developmentally key genes. For example, intrinsic fragility or biomechanical stress from mutations could activate regeneration programs that act as surveillance mechanisms. One prediction of this idea is that other genetic defects beyond *fras1* mutations should cause induction of regeneration-associated *fgf20a* expression.

To test if activation of regeneration-associated transcription is a general response to developmental defects, we performed a targeted loss-of-function (LOF) screen in the *fgf20a:EGFP* enhancer trap background. We selected 15 zebrafish orthologues of human disease genes that, when mutated in humans, cause congenital syndromes involving the skin, skeleton, or craniofacial structures—tissues related to those where *fgf20a* is induced by experimental injury **(Fig. 3A)**. These included genes linked to Fraser syndromes (Fras1, Frem2, Grip1), osteogenesis imperfecta and related bone fragility (Bmp1a, Col1a1, Col1a2, Col2a1), skeletal dysplasia (Fgfr3, Flnb, Nek1, Ebp), limb and heart malformations (Tbx5a), and cutaneous or connective tissue disorders (St14, Lmna, Pex1, Aspa). We chose these genes based on documented human pathologies and evidence from zebrafish models: for instance, *fras1*/*pinfin* and *frem2*/*blasen* mutants present fin blistering^31^, *bmp1a* mutants have ‘frilly fins’ owing to deficient collagen processing^32^, *tbx5a*/*heartstrings* mutants exhibit pectoral fin and heart malformations^33^, zebrafish *fgfr3* mutants show craniofacial malformations and delayed ossification similar to human skeletal dysplasia^34^, and CRISPR-created *imna* mutants present muscle and mobility defects analogous to muscular laminopathies^35^.

**Fig. 3.**
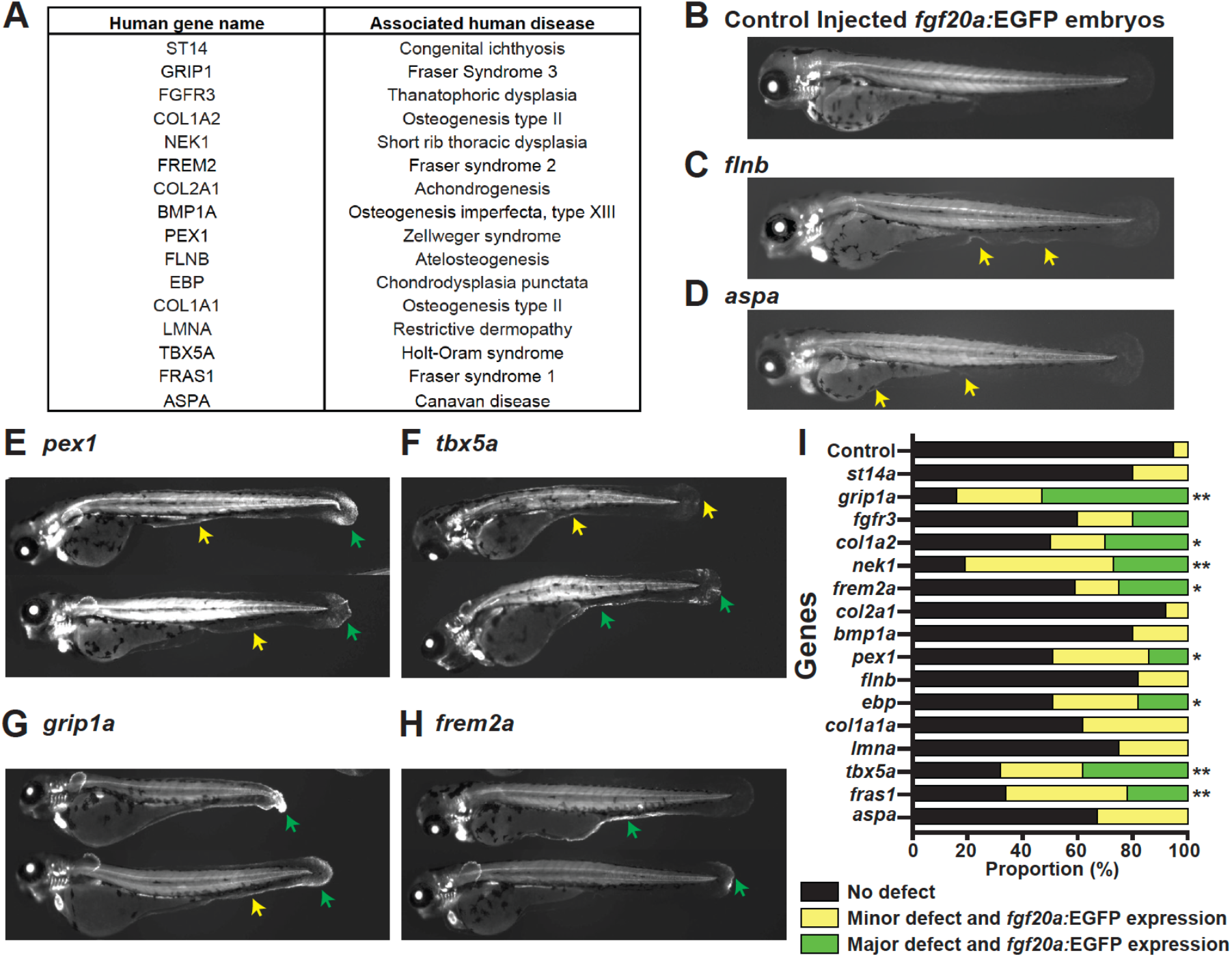
Loss of function mutations in zebrafish orthologues of human developmental disorder associated genes display degenerative phenotypes and activated *fgf20a* expression. **(A)** List of human genes included in the CRISPR-Cas9 F_0_ screen in the *fgf20a:EGFP* enhancer trap background. **(B-H)** Representative images from screen for zebrafish orthologues of human genes in **(A)**, as well as non-targeting control **(B). (I)** Quantification of phenotypes observed in the screen. Each larva was classified into either of 3 categories, namely, No expression, Minor or major defects as well as *fgf20a:*EGFP trap fluorescence (n > 12). Statistical significance was tested using Fisher’s exact test with correction for multiple comparisons. *p<0.05, *p<0.01.

Injection of CRISPR/Cas9 reagents targeting these genes resulted in larvae with varying degrees of morphological defects, most prominently in the fin fold and craniofacial tissues. In nearly every case where tissue integrity was visibly compromised, we observed ectopic activation of the *fgf20a:EGFP* enhancer trap in the affected regions **(Fig. 3B–H)**. The induced GFP expression domains extended beyond normal developmental patterns of *fgf20a*, which are typically restricted to the fin mesenchyme and various cranial structures, and instead appeared broadly in degenerating fin fold epithelia and malformed skeletal elements. Control-injected embryos, by contrast, generally exhibited *fgf20a:EGFP* expression in its canonical developmental domains, with no ectopic activity. Statistical analyses indicated that approximately half of the tested genes produced a significant enrichment of larvae in the “defect plus ectopic *fgf20a* expression” category **(Fig. 3I)**. Together, our experiments indicate that mutation-induced disruption of epithelial integrity, skeletal fragility, or tissue morphogenesis are broadly sufficient to engage *fgf20a* TREEs outside of an experimental injury context.

### Ectopic activation of other known regeneration responses in *fras1* mutant larvae

A second prediction of our hypothesis is that loss of *fras1* function should trigger regeneration-responsive enhancers in addition to those linked to *fgf20a*. To test this prediction, we focused on two well-characterized TREEs: *LEN*, which is required for regeneration-associated expression of the *leptin b* gene and directs expression of reporter transgenes in regenerating fin folds, adult fins, and adult hearts^3^; and *REN*, which is linked to the *runx1* locus and directs expression of reporter transgenes in injured and regenerating heart **(Fig. 4A)**^36^. As *LEN* directs expression in various tissues including both fin and heart, we tested whether REN also acts as a TREE in different tissues and directs expression in regenerating finfold. Indeed, we detected regeneration-responsive expression in mesenchyme and notochord of *RENcfos:EGFP* larvae. In wild-type animals, neither TREE directs expression of a reporter in intact fin folds, but they both direct local expression upon amputation of fin folds (**Fig. 4B**)^3,36^. To directly test whether inherited disorder of tissue degeneration is sufficient to engage TREEs, we crossed *LENP2:EGFP* or *RENcfos:EGFP* reporter transgenes into the *fras1* background. Strikingly, *fras1* mutants exhibited robust, ectopic activation of both reporters in degenerating finfold tissue by 4 dpf, in patterns closely resembling their normal injury-induced expression domains **(Fig. 4C and 4D)**. This response was highly penetrant, observed in nearly all mutant larvae examined, while control siblings showed no detectable reporter expression in intact fin folds. Thus, the *fras1* mutation does not only alter *fgf20a* regulation but engages at least two additional independent TREEs, supporting the view that complex regeneration programs are engaged in the context of developmental disorders.

**Fig. 4.**
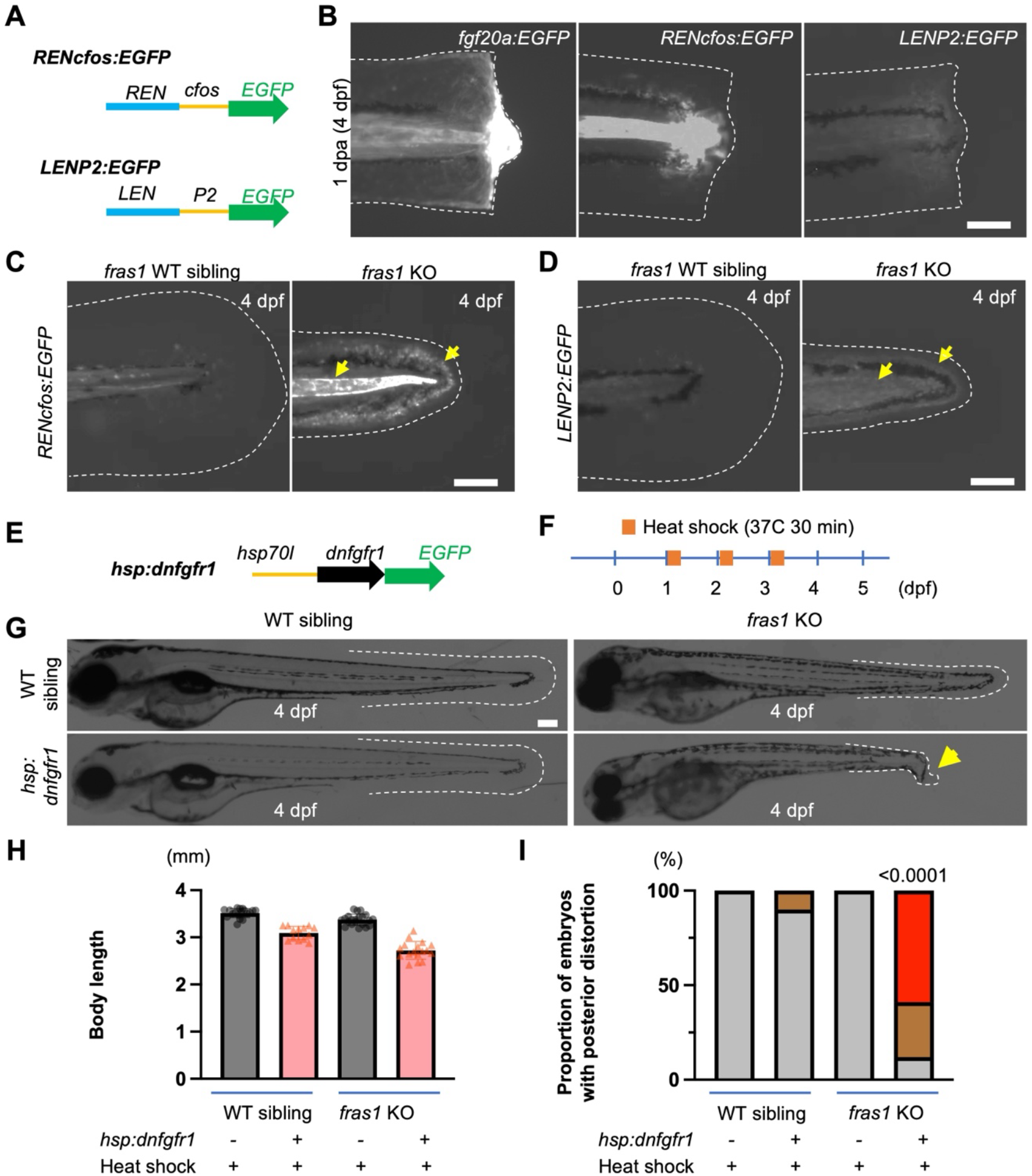
*fras1* phenotypes activate additional TREEs and are exacerbated by Fgf receptor blockade. **(A)** TREE reporter lines with *runx1* linked enhancer (*REN*) and *leptin b* linked enhancer (*LEN*) upstream of minimal promoter-EGFP cassettes. *cfos*, a 100 bp promoter of mouse *cfos* gene. *P2*, a 2 kb promoter of zebrafish *leptin b*. **(B)** Expression of reporters, *fgf20a:EGFP, RENcfos:EGFP* and *LENP2:EGFP* in regenerating larval fins at 4 dpf after fin amputation at 3 dpf. *fgf20a:EGFP* has regeneration-responsive expression in epithelia, mesenchyme, notochord and muscle, while *RENcfos:EGFP* and *LENP2:EGFP*, have regeneration-responsive expression in notochord and mesenchyme. **(C)** Skin degeneration by *fras1* mutation induces EGFP expression in notochord and mesenchyme (yellow arrows) of *RENcfos:EGFP* larvae. **(D)** Skin degeneration by *fras1* KO induces EGFP expression in mesenchyme (yellow arrows) of *LENP2:EGFP* larvae. **(E)** A heat shock-inducible dominant negative Fgf receptor line. **(F)** Heat shock is induced by incubating larvae in 37°C prewarmed eggwater for 30 min repeatedly at 1, 2, and 3 dpf. **(G)** Representative larvae of *hsp*:*dnfgfr1* and *fras1* mutant line and their negative controls at 4 dpf after three repeated heat shocks at 37°C for 30 min. Skin degeneration in *fras1* mutants does not activate *hsp70l* promoter without heat shock. **(H and I)** Bar graphs of body length and proportions of the larvae from **(G)**. Blocking Fgf signaling slightly shortens larval body length of larvae, but *fras1* mutations sensitize the posterior distortion.

### Genetic tempering of Fgf receptor signaling exacerbates *fras1* degenerative phenotypes

While evidence to this point indicates that genetic mutations defects can lead to regenerative responses in developing embryos, it was unclear if these responses provide function. That is, whether regeneration programs have protective effects against tissue degeneration caused by developmental defects. Because *fgf20a* is ectopically induced in *fras1* mutants, we reasoned that Fgf signaling might be part of a compensatory response that helps tissues tolerate degeneration. We therefore tested both directions of pathway perturbation. To assess whether increasing Fgf signaling could ameliorate the phenotype (and potentially extend survival, as most *fras1* mutants die within ∼2 weeks), we attempted localized ligand overexpression. Ubiquitous Fgf ligand delivery is known to cause early patterning defects such as dorsalization^37^, so we used *itgb4* regulatory sequences — reported to drive expression in larval fin fold epithelia^38^ — to confine *fgf20a* overexpression to the tissue of interest. In parallel, to test whether reducing Fgf signaling unmasks or worsens degeneration, we crossed *fras1* mutants to *hsp70l:dnfgfr1-EGFP* animals and induced the dominant-negative receptor by heat shock at larval stages **(Fig. 4E and 4F)**.

Targeted *fgf20a* overexpression under *itgb4* control was not tolerated, producing embryonic lethality in F_0_ animals all of which were arrested at epiboly stages and precluding assessment of improved responses — this is consistent with a narrow tolerance to elevated Fgf dosage even when spatially restricted. In contrast, temporally controlled Fgfr blockade yielded clear and interpretable outcomes. Induction of dnFgfr1 produced the expected general growth impairment (shortened body length) in wild-type and heterozygous controls and exacerbated fin fold degeneration in *fras1* mutants. Notably, a higher proportion with posterior body distortion than their heat-shocked siblings lacking the mutation **(Fig. 4G - 4I)**. Together, these results indicate that endogenous Fgf signaling is protective with respect to *fras1* phenotypes, preserving tissue integrity in the face of genetic mutations.

## DISCUSSION

Our findings suggest a broader role for regeneration programs than previously appreciated. In addition to being deployed for homeostatic maintenance and in response to acute injury, regeneration-associated enhancers and signaling pathways can also be engaged by early developmental defects. This raises the possibility that regeneration modules function not only to restore tissue after loss, but also to buffer the inherent fragility of morphogenesis. Such a buffering role could help explain why regeneration programs are preserved in vertebrates like zebrafish: by responding to a wide range of stresses—including genetic mutations, mechanical perturbations, or epithelial instability—these programs may contribute to developmental robustness and organismal survival. In this sense, regeneration is not simply a facultative repair response but a lifelong integrated component of tissue homeostasis and evolutionary fitness.

Several caveats temper this interpretation. The present work focuses on finfold epithelia and a small set of congenital disorder genes; it remains unclear how generalizable this buffering role is across other tissues, organs, or vertebrate lineages with limited regenerative ability. Moreover, our assays identify enhancer activation and phenotypic buffering but do not yet reveal the upstream sensors of tissue instability or the molecular logic by which developmental stress is distinguished. Future studies using single-cell transcriptomic and epigenomic approaches could clarify how regenerative circuits are rewired in genetic mutants, while cross-species comparisons may reveal whether the capacity to activate regeneration in response to developmental defects has been selectively maintained or diminished during evolution. Identifying these upstream signals and their integration with enhancer activity will be critical for testing if and how regeneration functions as a conserved developmental safeguard and whether this property can be leveraged therapeutically to ameliorate congenital disease.

The buffering role we suggest aligns with a broader concept in genetics: phenotypic variability. Many congenital mutations produce highly variable outcomes between individuals, even with identical genotypes. Such variability is thought to reflect compensatory mechanisms that mitigate developmental defects^39^. Regeneration programs, deployed through enhancer activation, may represent one such buffering system, limiting the impact of genetic lesions on tissue morphology. By connecting regeneration biology with developmental disease and variability, this work supports a model in which regeneration programs contribute to tissue robustness, and highlights enhancer elements as a molecular interface between genetic lesions and organismal outcome.

## MATERIALS AND METHODS

### Zebrafish maintenance and husbandry

Adult zebrafish (Danio rerio) of the outbred Ekkwill (EK) strain, up to 18 months of age, were maintained under standard conditions. Fish were housed in a recirculating system on a 14 h light/10 h dark cycle and fed twice daily with a combination of live brine shrimp and commercial flake food. Water temperature was maintained at 26–28 °C, conductivity at 500–700 µS, and pH at 7.0–7.5. Wild-type or transgenic animals, including the *fgf20a* enhancer trap line (allele number *HGn21A*) and the *hsp70:dnfgfr1-egfp* line (allele number *pd1)* were used as indicated^27,40^. All experiments were approved by and performed in accordance with institutional animal care and use protocols at Duke University and the University of Wisconsin–Madison.

### ENU mutagenesis in *fgf20a:EGFP* reporter fish

ENU mutagenesis was performed essentially as described^41,42^. Homozygous *fgf20a:EGFP* males (11 fish, ∼9 months old) were treated weekly with 3.3 mM ENU for 1 h over six consecutive weeks in fish water containing 10 mM sodium phosphate buffer (pH 6.5). To assess mutation efficiency, treated males were test-crossed to females homozygous for recessive pigmentation mutations (*slc45a2*^*b4/b4*^ or *kita*^*b5/b5*^) which is easily scored by 3 dpf ^43^. The ratio of pigmentation mutants was 4 in 1323 (*slc45a2*^*b4/*^*\**) and 4 in 962 (*kita*^*b5/*^*\**) which is high enough for the following mutant screen. Mutagenized males were mated to *fgf20a*:*EGFP* homozygous females and generated 11 F_1_ families (a total of 190 F_1_ fish). Intercrossing of the F_1_ fish generated 117 F_2_ families (a total of 920 F_2_ fish). Intercrossing of pairs of F_2_ fish in each family provided 460 clutches of F_3_ embryos. Twenty larvae per family underwent finfold amputation at 3 dpf, and regenerating fin folds were assessed at 4 dpf. Larvae with decreased or increased *fgf20a*:*EGFP* (or different EGFP patterns) at Mendelian ratios were candidates for mutations affecting *fgf20a* regulation. Putative mutant families were out-crossed again to either EK wild-types or homozygous *fgf20a*:*EGFP* fish to generate F_4_ heterozygotes, which were in-crossed to generate F_5_ families. F_5_ families were used for to confirm the phenotype.

### Anesthesia and larval finfold amputation

For experimental manipulations, larvae were anesthetized in 0.02% tricaine methanesulfonate (MS-222, Sigma, buffered to pH 7.0 with Tris). To perform larval finfold amputations, 3 days post-fertilization (dpf) larvae were placed on an agar-coated Petri dish under a dissecting microscope. Using a sterile no. 15 surgical scalpel, the finfold was transected at the proximal edge of the pigment gap adjacent to the circulatory loop of the caudal vein. Larvae were transferred to fresh embryo medium and allowed to recover at 28 °C. Regeneration was assessed at 1 dpa by monitoring gross morphology and EGFP expression.

### Imaging

Whole-mount larval images were acquired using a Zeiss AxioZoom V16 stereo fluorescence microscope equipped with GFP filter sets. Larvae were anesthetized in 0.02% Tricaine and mounted laterally on a 1% low-melting agarose bed in embryo medium for imaging. Brightfield and fluorescence images were collected using identical exposure settings across groups for comparability. Images were processed using Zen (Zeiss) and Fiji (ImageJ) software.

### Whole genome sequencing

For bulk WGS, *fras1* mutant carriers were in-crossed to produce clutches containing homozygous mutants and wild-type siblings. At 5 dpf, pools of ∼30 homozygous mutant larvae and ∼30 wild-type siblings were collected. Genomic DNA was extracted using the DNeasy Blood and Tissue Kit (Qiagen, Cat. 69504) according to manufacturer’s protocol. Libraries were prepared using the NEBNext Ultra II FS DNA Library Prep Kit (NEB, Cat. E7805), quantified with Qubit dsDNA HS Assay (ThermoFisher), and fragment sizes confirmed on a Bioanalyzer (Agilent). Sequencing was performed at BGI Genomics on a DNBseq platform with 150 bp paired-end reads, targeting at least 50 million read pairs per pool. Reads were trimmed with Trimmomatic, aligned to the Zebrafish GRCz11 reference genome using BWA-MEM, and variants were called with GATK HaplotypeCaller. Variants unique to mutant pools were identified, and filtering steps removed low-confidence calls, sequencing errors, and background polymorphisms from the EK strain. Homozygosity mapping identified a nonsense variant in fras1.

### Gene disruption by CRISPR/Cas9

CRISPR single-guide RNA (sgRNA) target sites were designed using the chopchop web tool (https://chopchop.cbu.uib.no/). Genomic DNA sequences retrieved from Ensembl GRCz10 or z11 (https://useast.ensembl.org/Danio_rerio/Info/Index) were used for the target site searches. Target sequences were selected that had no predicted off-target sites in the reference genome and no mismatches at loci to be targeted based on the whole-genome sequencing data of the laboratory EK strain. Target-specific Alt-R crRNA and common Alt-R tracrRNA were synthesized by IDT, and each RNA was dissolved in duplex buffer (IDT) as 1 mg/uL stock solution. Stock solutions were stored at -80 C. To prepare the crRNA:tracrRNA duplex, equal volumes of 1 mg/uL Alt-R crRNA and 1 mg/uL Alt-R tracrRNA stock solutions were mixed together and annealed by heating in a PCR machine: 95 C, 5 min; followed by gradual cooling on the bench for a few minutes. The 1 mg/mL crRNA:tracrRNA duplex stock solution was mixed with equal volume of 1 mg/mL Cas9 protein (PNA BIO), incubated at 37 C for 5 min and kept on ice prior to injection. The gRNA target sequences are listed in SI Appendix, Table S1.

### Blocking Fgf signaling by genetically inducing dominant-negative *fgfr1*

Embryos from crossing *hsp70l*:*dnfgfr1*-*EGFP*^+/-^;*fras1*^*/+^ and *fras1*^*/+^ fish were all heat shocked by transferring them into pre-warmed egg water for 30 min at 37°C and then transferred into egg water at room temperature. Heat shock was performed at 1, 2 and 3 dpf and the phenotype changes were assessed at 4 dpf. Anesthetized larvae were imaged using a Zeiss AxioZoom microscope at the time points indicated. All raw images were processed using either ZEN (Zeiss) or FIJI software. Lengths of larval bodies were measured using FIJI software as a longest straight line spanning the body projection. Proportions of embryos with posterior distortion after blocking Fgf signaling were calculated based on the numbers of embryos without/with mild/severe phenotype at the posterior region and analyzed using Fisher’s exact test.

All guide RNA targeting sequence information are available in SI Appendix, Table S1.

## Data and Materials Availability

New sequencing data have been deposited in NCBI under project accession code PRJNA1329955.

## Acknowledgements

We thank Morgridge Institute zebrafish facility staff and Duke University zebrafish facility staff for zebrafish care; Koichi Kawakami for transgenic *fgf20a*:EGFP enhancer trap animals; Atsushi Kawakami for *itgp4* plasmid; Colin Doran, Helen Rueckert, Tuyet Nguyen, and Sophie Dornbaum for contributions to experiments; and K.D.P. laboratory members for comments on the manuscript.

## Author contributions

Conceptualization, K.A., S.B., and K.D.P.; methodology, K.A., S.B., and K.D.P.; formal analysis, K.A., S.B., and K.D.P.; investigation, K.A., S.B., J.W., J.D.T., and K.D.P.; writing – original draft, K.A. and S.B.; writing – review & editing, K.A., S.B., and K.D.P.; resources and funding acquisition, K.D.P. All authors read and commented on or edited the manuscript.

## Supplementary Information

**Fig S1.**
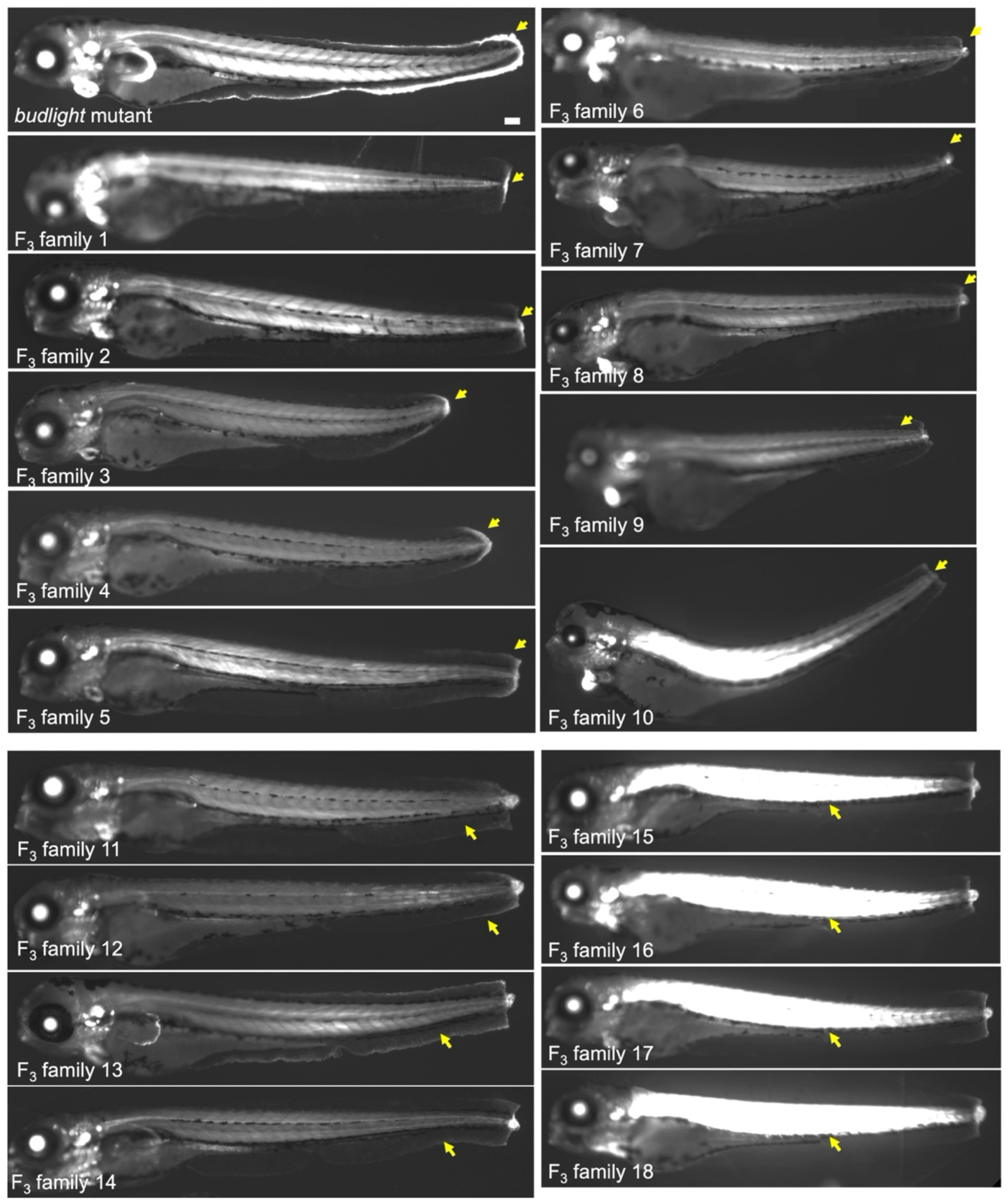
Phenotypes of F_3_ larvae. Ten F_3_ families contain larvae with limited regeneration phenotypes (yellow arrows) associated with EGFP expression, in both 1^st^ and 2^nd^ repeated screenings. F_3_ family 1-5 displayed reduced size of regenerating tissues, and F_3_ family 5-9 displayed EGFP expression at the edge of finfold. One of the mutants (named “*budlight*”) exhibited a strong regeneration defect linked to severely degenerated finfold in the absence of injury. Scale bar is 100 μm.

**Fig S2.**
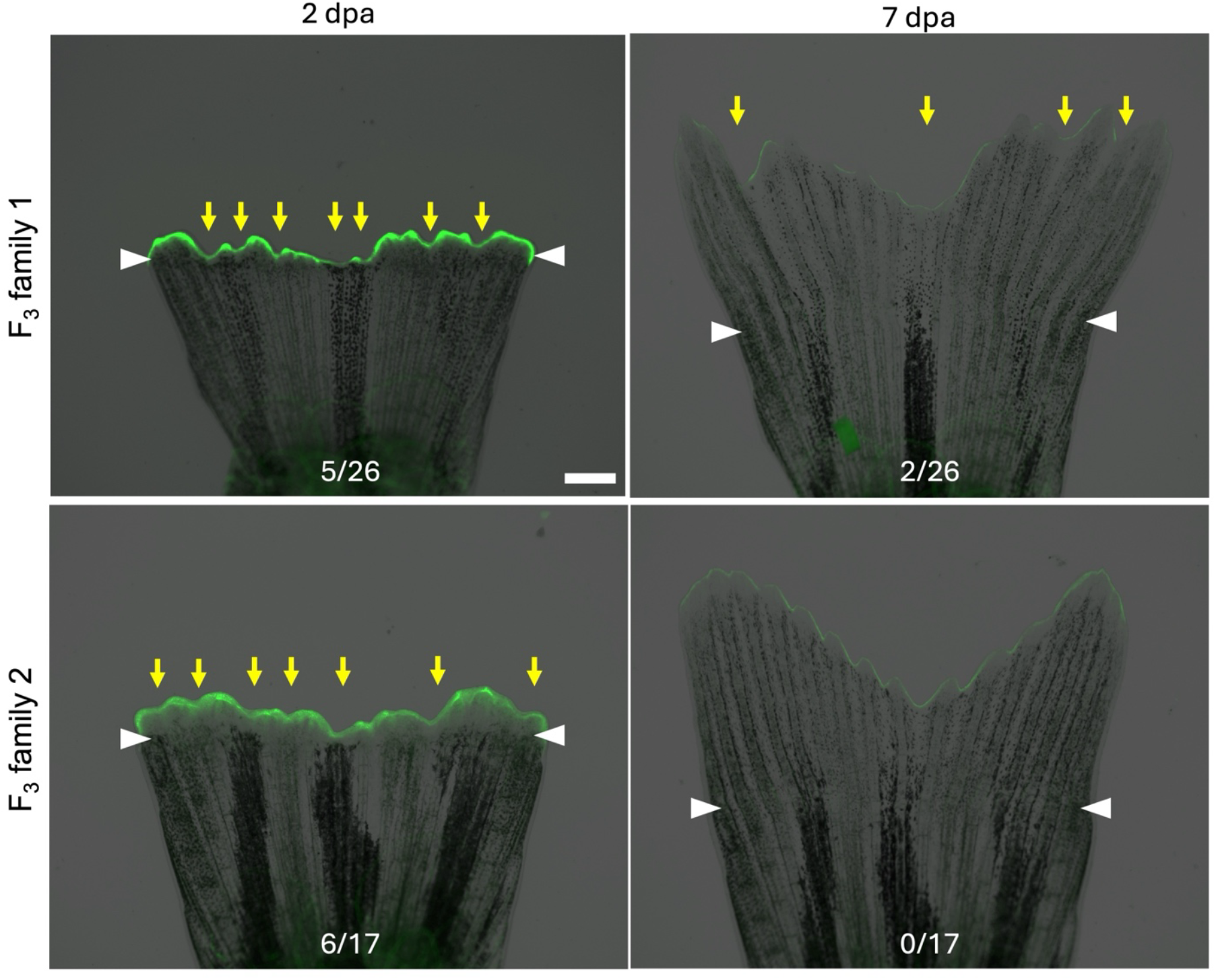
F_3_ adults with limited regenerating tissues. Two families exhibiting a regeneration phenotype in Fig. S1A contain adult fish with abnormal regeneration (Yellow arrows) at 2 dpa during caudal fin regeneration (left panels), although fin regeneration was grossly normal at 7 dpa (right panels). Ratios representing abnormal regeneration among total family members are at bottom of images. White arrow heads indicate amputation planes. Scale bar is 1 mm.

**SI Appendix, Table S1**.

**Table S1 lists the guide RNA targeting sequences (Length, 20 nt) used for the F**_**0**_ **screen of zebrafish orthologues of human congenital defect associated genes. For all genes, a combination of 2-3 guide RNAs in equimolar quantities was used**.

## REFERENCES

1. Poss, K.D. (2010). Advances in understanding tissue regenerative capacity and mechanisms in animals. Nat. Rev. Genet. 11, 710–722. 10.1038/nrg2879.

2. Poss, K.D., and Tanaka, E.M. (2024). Hallmarks of regeneration. Cell Stem Cell 31, 1244–1261. 10.1016/j.stem.2024.07.007.

3. Kang, J., Hu, J., Karra, R., Dickson, A.L., Tornini, V.A., Nachtrab, G., Gemberling, M., Goldman, J.A., Black, B.L., and Poss, K.D. (2016). Modulation of tissue repair by regeneration enhancer elements. Nature 532, 201–206. 10.1038/nature17644.

4. Goldman, J.A., and Poss, K.D. (2020). Gene regulatory programmes of tissue regeneration. Nat Rev Genet 21, 511–525. 10.1038/s41576-020-0239-7.

5. Whitehead, G.G., Makino, S., Lien, C.-L., and Keating, M.T. (2005). fgf20 Is Essential for Initiating Zebrafish Fin Regeneration. Science 310, 1957–1960. 10.1126/science.1117637.

6. Shibata, E., Yokota, Y., Horita, N., Kudo, A., Abe, G., Kawakami, K., and Kawakami, A. (2016). Fgf signalling controls diverse aspects of fin regeneration. Development 143, 2920–2929. 10.1242/dev.140699.

7. Thompson, J.D., Ou, J., Lee, N., Shin, K., Cigliola, V., Song, L., Crawford, G.E., Kang, J., and Poss, K.D. (2020). Identification and requirements of enhancers that direct gene expression during zebrafish fin regeneration. Development 147, dev191262. 10.1242/dev.191262.

8. Wang, W., Hu, C.-K., Zeng, A., Alegre, D., Hu, D., Gotting, K., Granillo, A.O., Wang, Y., Robb, S., Schnittker, R., et al. (2020). Changes in regeneration-responsive enhancers shape regenerative capacities in vertebrates. Science 369. 10.1126/science.aaz3090.

9. Chen, Y., Hou, Y., Zeng, Q., Wang, I., Shang, M., Shin, K., Hemauer, C., Xing, X., Kang, J., Zhao, G., et al. (2025). Common and specific gene regulatory programs in zebrafish caudal fin regeneration at single-cell resolution. Genome Res. 35, 202–218. 10.1101/gr.279372.124.

10. Cigliola, V., Shoffner, A., Lee, N., Ou, J., Gonzalez, T.J., Hoque, J., Becker, C.J., Han, Y., Shen, G., Faw, T.D., et al. (2023). Spinal cord repair is modulated by the neurogenic factor Hb-egf under direction of a regeneration-associated enhancer. Nat. Commun. 14, 4857. 10.1038/s41467-023-40486-5.

11. Jimenez, E., Slevin, C.C., Song, W., Chen, Z., Frederickson, S.C., Gildea, D., Wu, W., Elkahloun, A.G., Ovcharenko, I., and Burgess, S.M. (2022). A regulatory network of Sox and Six transcription factors initiate a cell fate transformation during hearing regeneration in adult zebrafish. Cell Genom. 2, 100170. 10.1016/j.xgen.2022.100170.

12. Lin, W., Jia, X., Shi, X., He, Q., Zhang, P., Zhang, X., Zhang, L., Wu, M., Ren, T., Liu, Y., et al. (2025). Reactivation of mammalian regeneration by turning on an evolutionarily disabled genetic switch. Science 388, eadp0176. 10.1126/science.adp0176.

13. Shi, T., Kim, Y., Llamas, J., Wang, X., Fabian, P., Lozito, T.P., Segil, N., Gnedeva, K., and Crump, J.G. (2024). Long-range Atoh1 enhancers maintain competency for hair cell regeneration in the inner ear. Proc. Natl. Acad. Sci. 121, e2418098121. 10.1073/pnas.2418098121.

14. Niethammer, P., Grabher, C., Look, A.T., and Mitchison, T.J. (2009). A tissue-scale gradient of hydrogen peroxide mediates rapid wound detection in zebrafish. Nature 459, 996–999. 10.1038/nature08119.

15. Godwin, J.W., Pinto, A.R., and Rosenthal, N.A. (2013). Macrophages are required for adult salamander limb regeneration. Proc. Natl. Acad. Sci. 110, 9415–9420. 10.1073/pnas.1300290110.

16. Petrie, T.A., Strand, N.S., Yang, C.-T., Tsung-Yang, C., Rabinowitz, J.S., and Moon, R.T. (2014). Macrophages modulate adult zebrafish tail fin regeneration. Development 141, 2581–2591. 10.1242/dev.098459.

17. Jopling, C., Suñé, G., Faucherre, A., Fabregat, C., and Belmonte, J.C.I. (2012). Hypoxia Induces Myocardial Regeneration in Zebrafish. Circulation 126, 3017–3027. 10.1161/circulationaha.112.107888.

18. Zhao, Y., Xiong, W., Li, C., Zhao, R., Lu, H., Song, S., Zhou, Y., Hu, Y., Shi, B., and Ge, J. (2023). Hypoxia-induced signaling in the cardiovascular system: pathogenesis and therapeutic targets. Signal Transduct. Target. Ther. 8, 431. 10.1038/s41392-023-01652-9.

19. Hirose, K., Payumo, A.Y., Cutie, S., Hoang, A., Zhang, H., Guyot, R., Lunn, D., Bigley, R.B., Yu, H., Wang, J., et al. (2019). Evidence for hormonal control of heart regenerative capacity during endothermy acquisition. Science 364, 184–188. 10.1126/science.aar2038.

20. Cheung, M.Y., Jiang, C., Hassan, I.U., Wang, H., Guo, D., Dio, D.W., Yan, H., Sun, J., Qi, X., Cai, D., et al. (2025). Knockout of thyroid hormone receptor alpha a (thraa) enhances cardiac regeneration in zebrafish through metabolic and hypoxic regulation. Cell Commun. Signal. 23, 340. 10.1186/s12964-025-02350-5.

21. Ross, I., Omengan, D.B., Huang, G.N., and Payumo, A.Y. (2022). Thyroid hormone-dependent regulation of metabolism and heart regeneration. J. Endocrinol. 252, R71–R82. 10.1530/joe-21-0335.

22. Simões, M.G., Bensimon-Brito, A., Fonseca, M., Farinho, A., Valério, F., Sousa, S., Afonso, N., Kumar, A., and Jacinto, A. (2014). Denervation impairs regeneration of amputated zebrafish fins. BMC Dev. Biol. 14, 49. 10.1186/s12861-014-0049-2.

23. Dagenais, P., Blanchoud, S., Pury, D., Pfefferli, C., Aegerter-Wilmsen, T., Aegerter, C.M., and Jaźwińska, A. (2021). Hydrodynamic stress and phenotypic plasticity of the zebrafish regenerating fin. J. Exp. Biol. 224. 10.1242/jeb.242309.

24. Gupta, V., Gemberling, M., Karra, R., Rosenfeld, G.E., Evans, T., and Poss, K.D. (2013). An Injury-Responsive Gata4 Program Shapes the Zebrafish Cardiac Ventricle. Curr. Biol. 23, 1221–1227. 10.1016/j.cub.2013.05.028.

25. Lin, C., Yao, E., Zhang, K., Jiang, X., Croll, S., Thompson-Peer, K., and Chuang, P.-T. (2017). YAP is essential for mechanical force production and epithelial cell proliferation during lung branching morphogenesis. eLife 6, e21130. 10.7554/elife.21130.

26. López-Anguita, N., Gassaloglu, S.I., Stötzel, M., Bolondi, A., Conkar, D., Typou, M., Buschow, R., Veenvliet, J.V., and Bulut-Karslioglu, A. (2022). Hypoxia induces an early primitive streak signature, enhancing spontaneous elongation and lineage representation in gastruloids. Development 149. 10.1242/dev.200679.

27. Nagayoshi, S., Hayashi, E., Abe, G., Osato, N., Asakawa, K., Urasaki, A., Horikawa, K., Ikeo, K., Takeda, H., and Kawakami, K. (2007). Insertional mutagenesis by the Tol2 transposon-mediated enhancer trap approach generated mutations in two developmental genes: tcf7and synembryn-like. Development 135, 159–169. 10.1242/dev.009050.

28. Shibata, E., Ando, K., Murase, E., and Kawakami, A. (2018). Heterogeneous fates and dynamic rearrangement of regenerative epidermis-derived cells during zebrafish fin regeneration. Development 145, dev162016. 10.1242/dev.162016.

29. McGregor, L., Makela, V., Darling, S.M., Vrontou, S., Chalepakis, G., Roberts, C., Smart, N., Rutland, P., Prescott, N., Hopkins, J., et al. (2003). Fraser syndrome and mouse blebbed phenotype caused by mutations in FRAS1/Fras1 encoding a putative extracellular matrix protein. Nat. Genet. 34, 203–208. 10.1038/ng1142.

30. Talbot, J.C., Walker, M.B., Carney, T.J., Huycke, T.R., Yan, Y.-L., BreMiller, R.A., Gai, L., DeLaurier, A., Postlethwait, J.H., Hammerschmidt, M., et al. (2012). fras1 shapes endodermal pouch 1 and stabilizes zebrafish pharyngeal skeletal development. Development 139, 2804–2813. 10.1242/dev.074906.

31. Carney, T.J., Feitosa, N.M., Sonntag, C., Slanchev, K., Kluger, J., Kiyozumi, D., Gebauer, J.M., Talbot, J.C., Kimmel, C.B., Sekiguchi, K., et al. (2010). Genetic Analysis of Fin Development in Zebrafish Identifies Furin and Hemicentin1 as Potential Novel Fraser Syndrome Disease Genes. PLoS Genet. 6, e1000907. 10.1371/journal.pgen.1000907.

32. Asharani, P.V., Keupp, K., Semler, O., Wang, W., Li, Y., Thiele, H., Yigit, G., Pohl, E., Becker, J., Frommolt, P., et al. (2012). Attenuated BMP1 Function Compromises Osteogenesis, Leading to Bone Fragility in Humans and Zebrafish. Am. J. Hum. Genet. 90, 661–674. 10.1016/j.ajhg.2012.02.026.

33. Garrity, D.M., Childs, S., and Fishman, M.C. (2002). The heartstrings mutation in zebrafish causes heart/fin Tbx5 deficiency syndrome. Development 129, 4635–4645. 10.1242/dev.129.19.4635.

34. Sun, X., Zhang, R., Chen, H., Du, X., Chen, S., Huang, J., Liu, M., Xu, M., Luo, F., Jin, M., et al. (2020). Fgfr3 mutation disrupts chondrogenesis and bone ossification in zebrafish model mimicking CATSHL syndrome partially via enhanced Wnt/β-catenin signaling. Theranostics 10, 7111–7130. 10.7150/thno.45286.

35. Nicolas, H.A., Hua, K., Quigley, H., Ivare, J., Tesson, F., and Akimenko, M. (2022). A CRISPR/Cas9 zebrafish lamin A/C mutant model of muscular laminopathy. Dev. Dyn. 251, 645–661. 10.1002/dvdy.427.

36. Goldman, J.A., Kuzu, G., Lee, N., Karasik, J., Gemberling, M., Foglia, M.J., Karra, R., Dickson, A.L., Sun, F., Tolstorukov, M.Y., et al. (2017). Resolving Heart Regeneration by Replacement Histone Profiling. Dev. Cell 40, 392-404.e5. 10.1016/j.devcel.2017.01.013.

37. Dorey, K., and Amaya, E. (2010). FGF signalling: diverse roles during early vertebrate embryogenesis. Development 137, 3731–3742. 10.1242/dev.037689.

38. Liu, Z., Meng, Y., Ishikura, A., and Kawakami, A. (2024). Live tracking of basal stem cells of the epidermis during growth, homeostasis, and injury response in zebrafish. Development 151. 10.1242/dev.202315.

39. Félix, M.-A., and Barkoulas, M. (2015). Pervasive robustness in biological systems. Nat. Rev. Genet. 16, 483–496. 10.1038/nrg3949.

40. Lee, Y., Grill, S., Sanchez, A., Murphy-Ryan, M., Poss, K.D. (2005). Fgf signaling instructs position-dependent growth rate during zebrafish fin regeneration. Development 132, 5173–5183. 10.1242/dev.02101.

41. Bruijn, E. de, Cuppen, E., and Feitsma, H. (2009). Zebrafish, Methods and Protocols. Methods Mol. Biol. 546, 3–12. 10.1007/978-1-60327-977-2_1.

42. Trevarrow, B. (2011). Techniques for optimizing the creation of mutations in zebrafish using N-ethyl-N-nitrosourea. Lab Anim. 40, 353–361. 10.1038/laban1111-353.

43. Riley, B.B., and Grunwald, D.J. (1995). Efficient induction of point mutations allowing recovery of specific locus mutations in zebrafish. Proc. Natl. Acad. Sci. USA. 92, 5997–6001. 10.1073/pnas.92.13.5997.

